# Larval growth rate affects wing shape more than eyespot size in the seasonally polyphenic butterfly *Melanitis leda*

**DOI:** 10.1101/2023.12.11.571078

**Authors:** Freerk Molleman, Elizabeth M. Moore, Sridhar Halali, Ullasa Kodandaramaiah, Dheeraj Halali, Erik van Bergen, Paul M. Brakefield, Vicencio Oostra

## Abstract

Butterflies often show adaptive phenotypic plasticity where environmental cues during early stages are used to produce a phenotype that maximizes fitness in the environment experienced by adults. Many tropical satyrine butterflies (Nymphalidae: Satyrinae) are seasonally polyphenic and produce distinct wet- and dry-season form adults providing tight environment-phenotype matching in seasonal environments. Dry-season forms, which are expressed in the dry season, can be induced in the laboratory by growing larvae at low temperatures or on poor food quality. Since both these factors also tend to reduce larval growth rate, larval growth rate may be an internal cue that translates the environmental cues into the expression of phenotypes. If this is the case, we predict that slower-growing larvae would be more likely to develop a dry-season phenotype. To test this hypothesis, we measured both larval growth rate and adult phenotype (eyespot size and wing shape) of individuals of the common evening brown butterfly (*Melanitis leda*), reared at various temperatures and on various host-plant species. We found that among treatments, larvae with lower growth rates (low temperature, particular host plants) were more likely to develop dry-season phenotypes (small eyespots, falcate wing tips), but within treatments, larval growth rate was mainly linked to wing shape, not eyespot size. These relationships tended to be stronger for males than females as males showed a wider range of eyespot sizes and wing shapes. Overall, only plasticity in wing shape appears to be (partly) mediated by larval growth, and in a sex-specific manner.

## INTRODUCTION

Many organisms are adapted to the seasonality of their environment using phenotypic plasticity to generate seasonal forms that fit the environment during the respective seasons (seasonal polyphenism, Tauber et al. 1986). Insects often show developmental plasticity where individuals use environmental cues during early stages to produce an adult phenotype that maximizes fitness in the environment experienced during the adult (Shapiro 1976; West-Eberhard 2003). These cues can for example be temperature, day length, or food quality (Yoon et al., 2023). For instance, some temperate polyphenic butterflies exhibit spring or summer phenotype depending on day length (Dorfmeister 1864; Rountree & Nijhout 1995). Moreover, species can use more than one cue, and these cues may interact with each other (Yoon et al. 2023). Parts of the physiological cascade that translates environmental cues into induction of the phenotype are well understood in some insect model organisms (Baudach & Vilcinskas 2021; Monteiro 2017; Oostra et al. 2011; Singh et al. 2020; Steward et al. 2022). However, we have limited insight into how multiple cues are integrated, and induce the expression of a suite of traits, especially in non-model organisms.

A prominent example of adaptive phenotypic plasticity in seasonal environments is seasonal polyphenism exhibited by many tropical satyrine (Nymphalidae: Satyrinae) butterflies (Brakefield & Larsen 1984; van Bergen & Oostra 2023). These butterflies express a wet-season form with large ventral eyespots along the wing margins, and a dry-season form with reduced eyespots and overall a cryptic wing pattern (Brakefield & Larsen 1984; Brakefield & Reitsma 1991). In addition to color pattern, a suite of life-history and behavioral traits covary in seasonal forms, including wing shape (Brakefield 1987). While most studies focus on the strong effects of temperature (Kooi & Brakefield 1999; Nokelainen et al. 2018; van Bergen et al. 2017; Windig 1992), other cues can interact with temperature affecting the eyespot size and overall wing pattern. For example, the quality of the host plant can interact with temperature in determining eyespot size or overall wing pattern (Kooi et al. 1996; Singh et al. 2020). Some parts of the cascade from cue to adult phenotype are well understood for the satyrine butterfly *Bicyclus anynana.* For example, we know how cues induce differences in the temporal expression of ecdysone hormone and gene expression, and how this affects wing pattern development (Oostra et al. 2011; Oostra et al. 2014; Tian & Monteiro 2022). However, it is mostly unknown how multiple environmental cues can drive wing pattern plasticity together, presumably by converging on the same regulatory system.

Larval growth rate could potentially integrate multiple cues for eyespot size in satyrine butterflies. Low temperatures and poor food quality experienced by larvae can induce the development of dry-season-form adults (small eyespots) in *Bicyclus* butterflies, and both of these factors also reduce larval growth rate (Kooi et al. 1996; Nokelainen et al. 2016; Windig 1992; Windig 1994). Even within a treatment, larval growth may be correlated with adult phenotype (Windig 1994). Therefore, it has been hypothesized that larval growth rate is part of the pathway that translates the environmental cues into the expression of phenotypes (Kooi et al. 1996; Windig 1992). However, it is not clear if there is a direct relationship between larval development and adult phenotype, or that each is correlated with these environmental factors, without a direct link between them. Moreover, these studies were limited to a small number of closely related species (mainly *B. anynana*) and usually focused on a single metric of adult phenotype, eyespot size.

Here we investigate how multiple cues affect the expression of seasonal polyphenism (in the common evening brown butterfly (*Melanitis led* L., Melanitiini, Satyrinae, Nymphalidae). *M. leda* is distantly related to *Bicyclus* species, and shows prominent polyphenism in eyespot size as well as wing shape (Brakefield 1987; Halali et al. 2019). We tested for the use of temperature and host plant as cues, and described provisional reaction norms for two populations. We predicted that *M. leda* would respond similarly to temperature and host plant: more wet-season phenotypes at higher temperatures and on better host plants. We also performed preliminary experiments to gauge interactions between temperature and humidity, expecting higher humidity to result in more wet-season phenotypes at a given temperature. We then studied the relationship between larval growth rate and two aspects of adult phenotype, eyespot size and wing shape. In three separate experiments, we manipulated larval growth rate by varying either only rearing temperature, both temperature and relative humidity, or host-plant species. If growth rate indeed mediates between environmental cues and adult phenotype, growth rate and phenotype are predicted to not only be related among treatments, but also within treatments (Figure 1).

**Figure 1.**
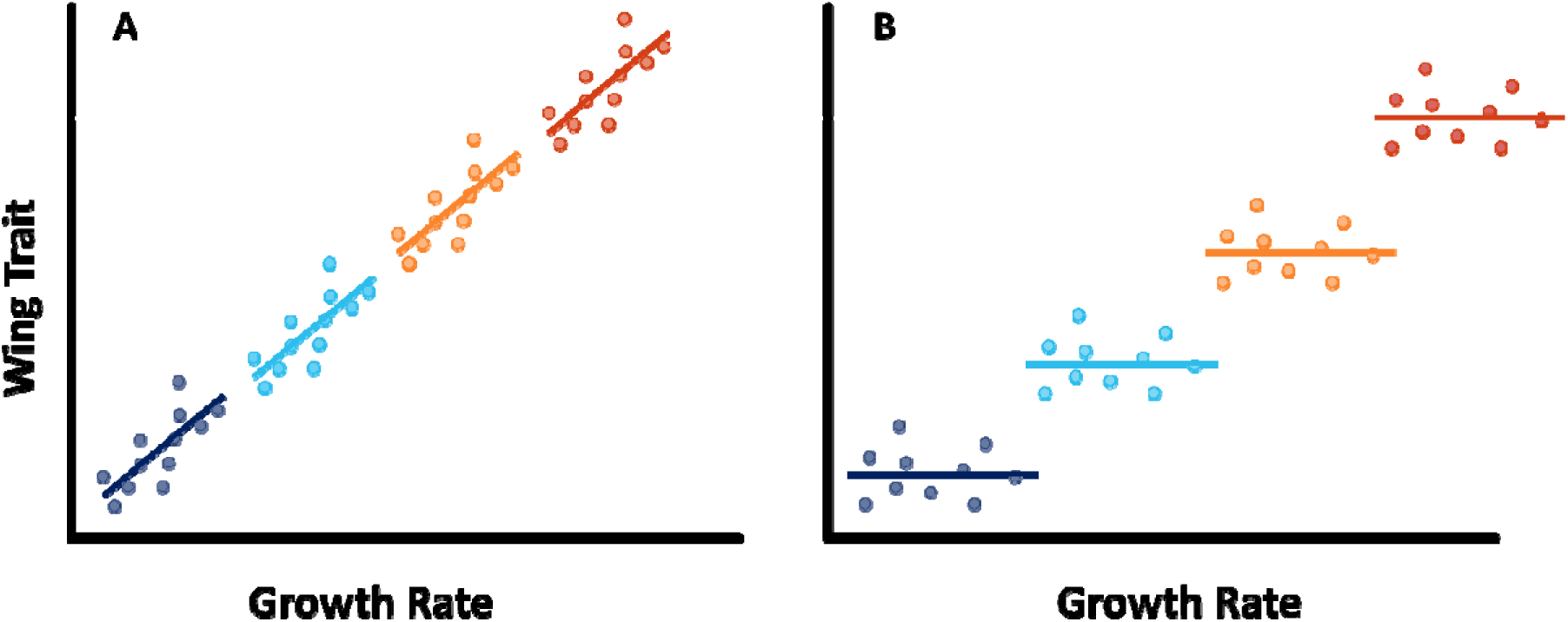
Predictions. A: If larval growth rate is part of the pathway that translates the environmental cues into the expression of phenotypes, we would expect to see a relationship between wing traits and growth rate across experimental treatment groups, as well as within treatments. B: If larval growth rate is not part of this pathway, we may still see a relationship between larval growth rate and adult phenotype among experimental treatment groups, but no such relationship within treatments.

## METHODS

### Model system

*M. leda* is among the most widespread butterflies, distributed from Africa to Asia and Oceania (Latorre 2018). It is able to use a wide variety of grass species (Poaceae) as host plant (Molleman et al. 2020b). *M. leda* displays seasonal phenotypic plasticity where wet-season forms have large and conspicuous eyespots, while dry-season forms have small eyespots (Fig. 2). Dry-season forms also have more falcate forewing tips and longer hindwing tails compared to the wet-season forms (Brakefield 1987; Halali et al. 2019). Field data suggest that this seasonal plasticity is to some extent mediated by temperature (Brakefield 1987; Halali et al. 2021; Roy et al. 2021). Furthermore, the variation among dry-season form wing patterns of *M. leda* represents one of the most multi-faceted polymorphisms in wing coloration found in any animal (Ruiter & Brakefield 1994).

**Figure 2.**
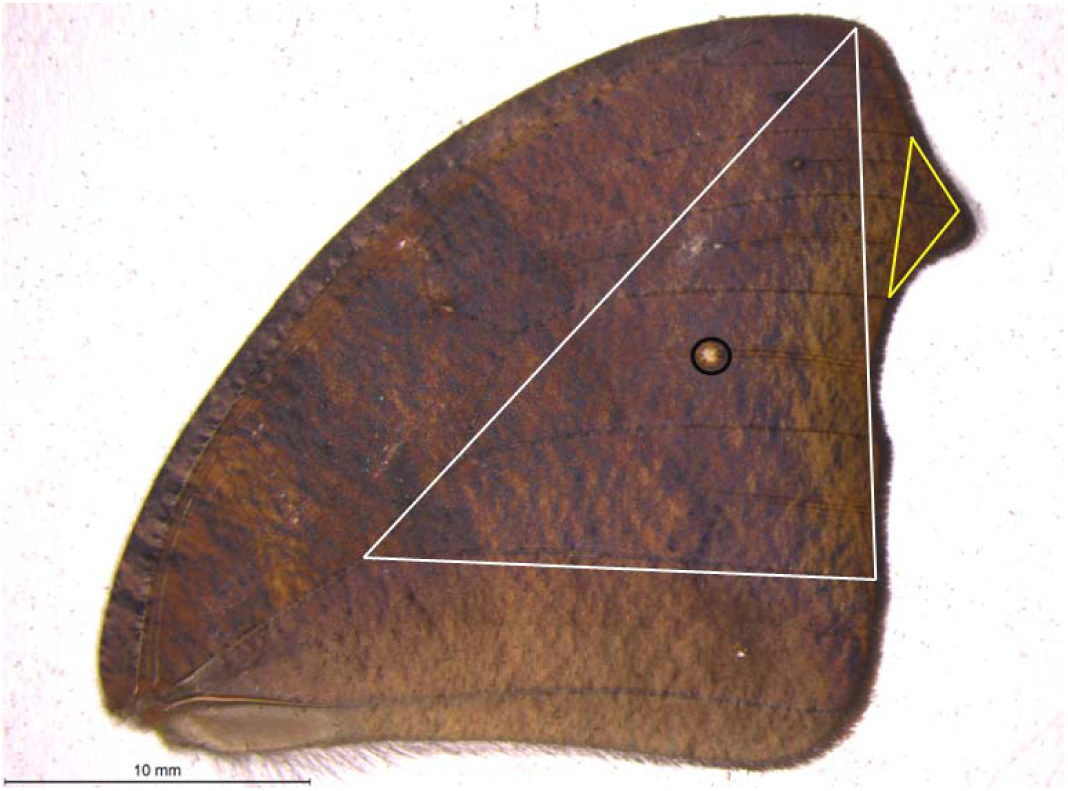
*M. leda* forewing with landmarks used in this study (female dry-season form from Ghanaian population). The white triangle denotes the proxy of wing area, the yellow triangle the proxy for wing shape, and the black circle eyespot size.

### General procedures and data collection

Three experiments were performed. The first experiment tested the effect of temperature on adult phenotype in a population from Ghana (Temperature Experiment). The second included effects of temperature and humidity using a population from South India (Temperature and Humidity Experiment). The third experiment tested for effect of host-plant species using the South-Indian population (Host-Plant Experiment). While some methodological details differ, the data taken from these experiments are comparable. In summary, caterpillars were reared on grasses growing in flower pots either from seeds or collected from the field (such as in Molleman et al. 2020b). The caterpillars were confined to the plants by placing the plants in fine-mesh sleeves. These sleeves were then kept in climate chambers that controlled temperature, and humidity, and had a 12/12 hour light schedule. Plants were watered every other day, and were replaced once the majority of leaf material had been consumed or general plant quality had deteriorated. Once larvae reached the 5^th^ instar, sleeves were inspected every 24 hours for prepupae and pupae. Prepupae and pupae were kept individually in cylindrical transparent containers of about 100ml until eclosion. Prepupae were left attached to the plant whenever possible: using adhesive tape, a piece of the plant where the prepupa was attached was fixed to the side or lid of the container, so that the prepupae could hang freely while pupating. Absorbent paper was placed at the bottom of the container. Hardened pupae were gently removed from the plant material to be weighed and sexed, and then placed back inside the container. Containers were kept upside down with the absorbent paper on the lid so that pupae and eclosing butterflies were easy to observe through the transparent bottoms. The sides of each of these containers were lined with a strip of paper on which the eclosing adult could climb to enable it to expand its wings. Containers were checked daily for adult eclosions. Butterflies were frozen one day after eclosion.

Data were collected on development time from egg hatching to pupation (larval development time), and from pupation to eclosion (pupal time). The eclosed butterflies were photographed or scanned and eyespots, forewing tip, and hindwing tails were measured using an ImageJ macro (Schindelin et al. 2015). For this study, we focused on the largest eyespot on the forewing to represent eyespot size and the forewing tip to represent wing shape (Figure 2).

### Temperature Experiment, Ghanaian population

We collected eggs from 14 females from a stock population established based on 80 females from Bobiri, Ghana. Sleeves contained 5-30 larvae with the average being ca. 20, and were fed ca. two-week-old corn seedlings. These sleeves were placed within environmental test chambers at 75% relative humidity. Larvae were reared at five temperatures; 19°C, 22°C, 25°C, 28°C and 31°C. Because of unexpected results, a follow-up experiment was carried out at three temperatures; 22°C, 25°C, and 28°C (results in Appendix 1). Photographs were taken using a Leica microscope imaging system.

### Temperature and Humidity Experiment, Indian population

Female *M. leda* butterflies were collected in Thiruvananthapuram, Kerala, South India, to provide eggs, and experiments were performed on three subsequent generations. Sleeves were set up with 15-17 caterpillars each, and were fed ca. two-week-old corn seedlings. Two incubators were used during subsequent experiments with combinations of four temperatures (19°C, 21°C, 27°C, and 31°C) and two levels of humidity (65% and 85% relative humidity). However, for technical reasons, a complete set of temperature and humidity combinations was not achieved so that the effect of humidity during rearing on adult phenotype could not be tested formally. The wings of the eclosed butterflies were scanned for phenotypic measurements (Konica Minolta, Bizhub 363).

### Host-Plant Experiments, Indian Population

The data presented here are based on the rearing of the fourth batch of larvae described in Molleman *et al*. (2020b). In short, female *M. leda* butterflies were collected in Thiruvananthapuram to provide eggs. Larvae were then grown on potted plants belonging to 18 species of grass in an incubator with a constant temperature and humidity of 24°C with 69% relative humidity. Photos of the wings of adults were taken with a Nikon D7000 camera under standard light inside a closed white styrofoam box with a shutter speed of 1/125 and an aperture of F14.

### Data analysis

Relative eyespot area was measured by dividing the area of the M3 eyespot by a proxy of total wing area (Figure 2). Wing shape was quantified as the surface of the wing-tip triangle divided by the proxy for wing area (Figure 2). Larval growth rate was calculated as (pupal mass)^1/3^ / (larval development time), following Tammaru & Esperk (2007). We first explored the distribution of growth rate, relative eyespot size, and wing shape for both sexes using density plots. While larval growth rate approached a normal distribution, relative eyespot size and wing shape were biased towards very small numbers and this could be corrected by using the square-root transformation. For ease of use, we use ‘eyespot size’ as a shorthand for ‘square root of relative eyespot size’ and ‘wing shape’ as a shorthand for ‘square root of relative size of the large forewing tip triangle’. Plots and preliminary statistical analyses revealed differences between the sexes in the relationship between growth rate, eyespot size, and wing shape, so we analysed males and females separately.

To generate provisional reaction norms for temperature, humidity, and host plants, the eyespot size and wing shape were plotted against treatment for all three experiments. For the Temperature and Humidity Experiment, data from a pilot experiment were included in which second instar larva were allocated to 85% RH at either 21°C or 27°C and sex of adults was not determined. For the Host-Plant Experiment, plants on which less than eight larvae were reared to adults were excluded, and plants were ordered according to average eyespot size or wing shape.

To test if growth rate predicts wing traits within treatments, we performed linear least square regression for each treatment separately. So, for each experiment and each treatment, we plotted a regression line and reported the statistics. All analyses were performed in R (R_Core_Team 2023) and graphs were made using the R package ggplot2 (Wickham 2016).

## RESULTS

### Cue use and reaction norms

Eyespot size had a bimodal distribution that appeared similar for all experiments and both sexes (Figures 3a & b). Eyespot size increased with increasing rearing temperature in both populations (Figures 3a & b). However, the relationship was more non-linear in the Ghanaian population with eyespot size being smaller when reared at 25°C than at 27°C (Figure 3a). Similar results were found in the follow-up experiment designed to verify this non-linear effect (See Appendix 1). Notably, the sexes showed similar temperature reaction norms for eyespot size (Figure 3a). At intermediate rearing temperatures, individuals with both wet- and dry-season form eyespots were produced, with a few intermediates (22°C, 25°C Figure 3a, 21°C, 27°C Figure 3b). Higher humidity appeared to result in more wet-season form individuals at intermediate temperatures (Figure 3b), but note that these are pilot data starting from 2^nd^ and 3^rd^ instar caterpillars, and these data are not included in further analyses. Average eyespot size also varied with host-plant species (Figure 3c). Males and females showed similar host-plant reaction norms for eyespot size (Figure 3c).

**Figure 3.**
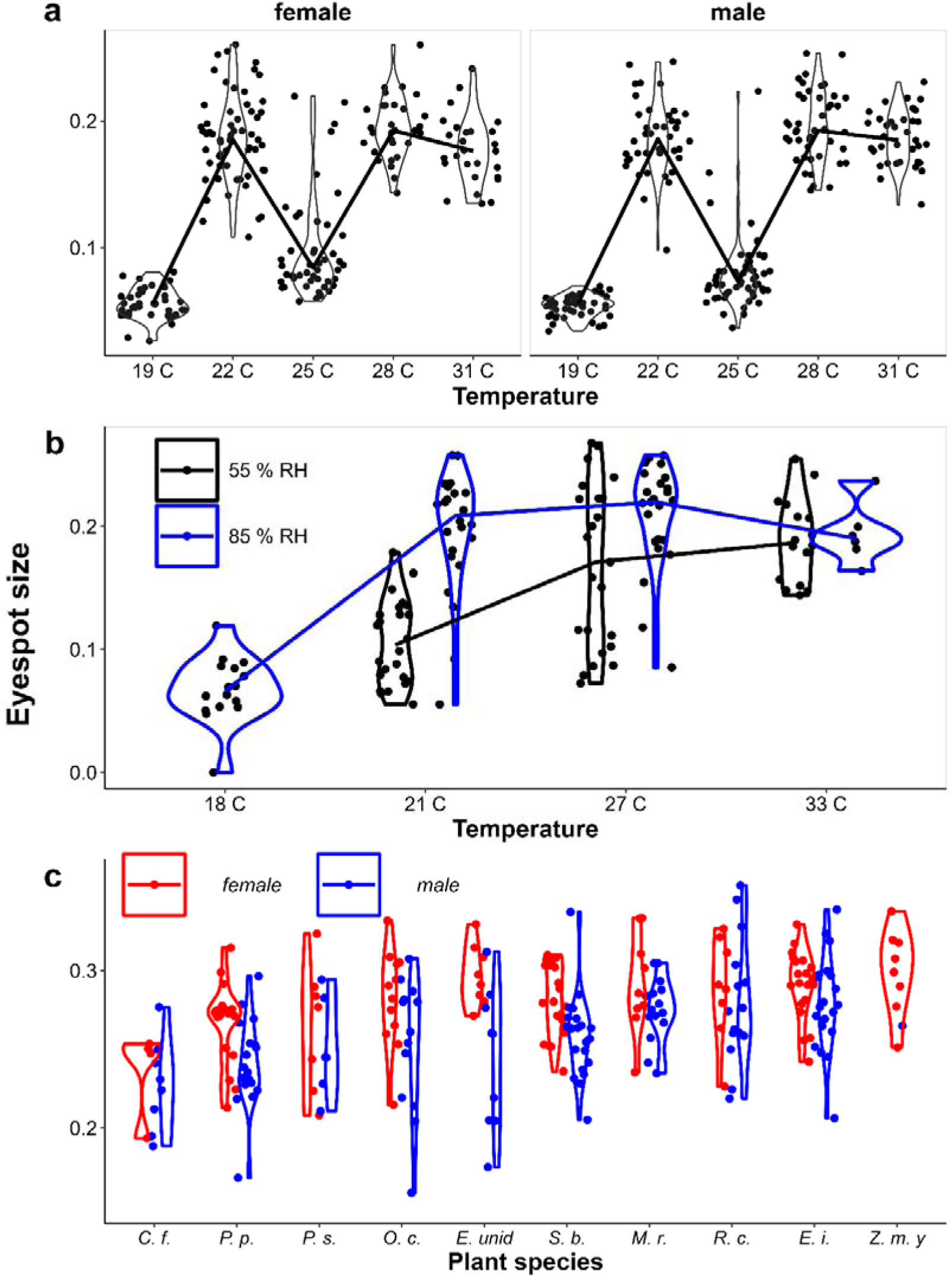
Provisional reaction norms of *M. leda* eyespot size for a) the Temperature Experiment, b) the Temperature and Humidity Experiment, and c) the Host-Plant Experiment. Note that for b the data for high humidity for temperatures 21° and 27°C are from a pilot experiment starting from 2^nd^ instar caterpillars and sex was not recorded. For 3c only plants with N > 7 were included and plants were sorted from small to large average eyespot size. No true dry-season form butterflies were produced in the Host-Plant Experiment so the y-axis was truncated. Full species names are given in Table 1.

**Table 1.** Results of linear regression for eyespot size. P-values of < 0.05 are in bold font with white background. Temp.= rearing temperature, Plant species are: *D. s. = Dendrocalamus strictus* (Roxb.) Nees, *A. c. = Axonopus compressus* (Sw.) P. Beauv, *O. unid =* Unidentified *Oplismenus* bush, *Z. m. o = Zea mays* L. mature plants, *Z. m. y = Zea mays* L. seedlings, *C. h. = Chrysopogon hackelii* (Hook.f.)*, B. mu. = Brachiaria mutica* (Forssk.) Stapf, *P. p. = Pennisetum polystachion* (L.) Schult., *C. d. = Cynodon dactylon* (L.) Pers., *P. s.= Paspalum scrobiculatum* L., *O. t. = Ochlandra travancorica* (Bedd.)*, O. c. = Oplismenus compositus* (L.), P. Beauv., *S. b. = Setaria barbata* (Lam.) Kunth, *B. mi. = Brachiaria milliformis. R. c. = Rottboellia cochinchinensis* (Lour.) Clayton, *C. f. = Cymbopogon flexuosus* (Nees ex St.) Watson, *E. unid =* Unidentified like *Eleusine, E. i. = Eleusine indica* (L.) Gaertn, *M. r.= Melinis repens* (Willd.) Zizka, *P. r. = Panicum repens* L., *T. a. = Triticum aestivum* L. N = the number of individuals reared to adult stage and measured, Estimate = the effect estimate of growth rate on eyespot size, S. E. = standard error of the estimate, Statistic = T or F statistic, p = p-value for the effect of growth rate of eyespot size.

| Exp. | Temp. | Humidity | Plant | sex | N | Estimate | S. E. | Statistic | p-value | R<sup>2</sup> | Adj. R<sup>2</sup> |
| --- | --- | --- | --- | --- | --- | --- | --- | --- | --- | --- | --- |
| Temperature Experiment, Ghanaian population |  |  |  |  |  |  |  |  |  |  |  |
|  | 19 C | 75% | Z. m. y | female | 39 | 0.90 | 0.99 | 0.91 | 0.37 | 0.02 | 0.00 |
|  | 22 C |  |  |  | 52 | 3.06 | 1.17 | 2.62 | **0.01** | 0.12 | 0.10 |
|  | 25 C |  |  |  | 37 | 0.75 | 1.41 | 0.54 | 0.60 | 0.01 | -0.02 |
|  | 28 C |  |  |  | 23 | 1.63 | 1.00 | 1.62 | 0.12 | 0.11 | 0.07 |
|  | 31 C |  |  |  | 25 | 0.62 | 1.61 | 0.39 | 0.70 | 0.01 | -0.04 |
|  | 19 C |  |  | male | 44 | 1.83 | 0.71 | 2.57 | **0.01** | 0.14 | 0.12 |
|  | 22 C |  |  |  | 32 | 0.89 | 1.51 | 0.59 | 0.56 | 0.01 | -0.02 |
|  | 25 C |  |  |  | 40 | -0.96 | 0.94 | -1.01 | 0.32 | 0.03 | 0.00 |
|  | 28 C |  |  |  | 41 | -0.78 | 1.00 | -0.78 | 0.44 | 0.02 | -0.01 |
|  | 31 C |  |  |  | 31 | -0.94 | 1.00 | -0.94 | 0.35 | 0.03 | 0.00 |
| Temperature and Humidity Experiment, Indian population |  |  |  |  |  |  |  |  |  |  |  |
|  | 18 C | 85% | Z. m. y | female | 8 | -3.78 | 6.19 | -0.61 | 0.56 | 0.06 | -0.10 |
|  | 21 C | 55% |  |  | 10 | 6.28 | 2.11 | 2.98 | **0.02** | 0.53 | 0.47 |
|  | 27 C | 55% |  |  | 14 | -3.21 | 8.37 | -0.38 | 0.71 | 0.01 | -0.07 |
|  | 33 C | 55% |  |  | 7 | 0.96 | 7.79 | 0.12 | 0.91 | 0.00 | -0.20 |
|  | 18 C | 85% |  | male | 8 | 1.80 | 4.35 | 0.41 | 0.69 | 0.03 | -0.13 |
|  | 21 C | 55% |  |  | 14 | 6.01 | 2.87 | 2.10 | 0.06 | 0.27 | 0.21 |
|  | 27 C | 55% |  |  | 8 | -11.87 | 11.41 | -1.04 | 0.34 | 0.15 | 0.01 |
|  | 33 C | 55% |  |  | 9 | -1.76 | 3.04 | -0.58 | 0.58 | 0.05 | -0.09 |
|  | 33 C | 85% |  |  | 3 | -12.29 | 29.43 | -0.42 | 0.75 | 0.15 | -0.70 |
| Host Plant Experiment, Indian population |  |  |  |  |  |  |  |  |  |  |  |
|  | 24 C | 69% | Z. m. y | female | 8 | -0.79 | 2.10 | -0.38 | 0.72 | 0.02 | -0.14 |
|  |  |  | P. p. |  | 18 | 0.44 | 1.08 | 0.41 | 0.69 | 0.01 | -0.05 |
|  |  |  | O. c. |  | 12 | -0.90 | 2.09 | -0.43 | 0.68 | 0.02 | -0.08 |
|  |  |  | S. b. |  | 19 | 0.43 | 1.35 | 0.32 | 0.75 | 0.01 | -0.05 |
|  |  |  | R. c. |  | 8 | -1.21 | 2.42 | -0.50 | 0.64 | 0.04 | -0.12 |
|  |  |  | E. unid |  | 8 | -2.49 | 2.86 | -0.87 | 0.42 | 0.11 | -0.04 |
|  |  |  | E. i. |  | 20 | 1.80 | 0.92 | 1.96 | 0.07 | 0.18 | 0.13 |
|  |  |  | M. r. |  | 10 | 3.66 | 2.50 | 1.46 | 0.18 | 0.21 | 0.11 |
|  |  |  | P. p. | male | 18 | 1.87 | 1.34 | 1.40 | 0.18 | 0.11 | 0.05 |
|  |  |  | O. c. |  | 13 | 1.48 | 2.02 | 0.73 | 0.48 | 0.05 | -0.04 |
|  |  |  | S. b. |  | 20 | 1.15 | 1.18 | 0.98 | 0.34 | 0.05 | 0.00 |
|  |  |  | R. c. |  | 15 | 3.13 | 1.42 | 2.21 | **0.05** | 0.27 | 0.22 |
|  |  |  | C. f. |  | 8 | 1.93 | 6.32 | 0.30 | 0.77 | 0.02 | -0.15 |
|  |  |  | E. unid |  | 9 | 18.51 | 9.21 | 2.01 | 0.08 | 0.37 | 0.28 |
|  |  |  | E. i. |  | 20 | 2.14 | 1.33 | 1.60 | 0.13 | 0.12 | 0.08 |
|  |  |  | M. r. |  | 15 | 1.06 | 1.21 | 0.87 | 0.40 | 0.06 | -0.02 |

In contrast to eyespot size, wing shape did not show a bimodal distribution, as a large number of individuals showed intermediate wing shapes (Figure 4). Otherwise, results for wing shape were similar to those for eyespot size: increasing temperature and humidity caused more wet-season phenotypes (less falcate wing tips), and there was an effect of host plant (Figure 4).

**Figure 4.**
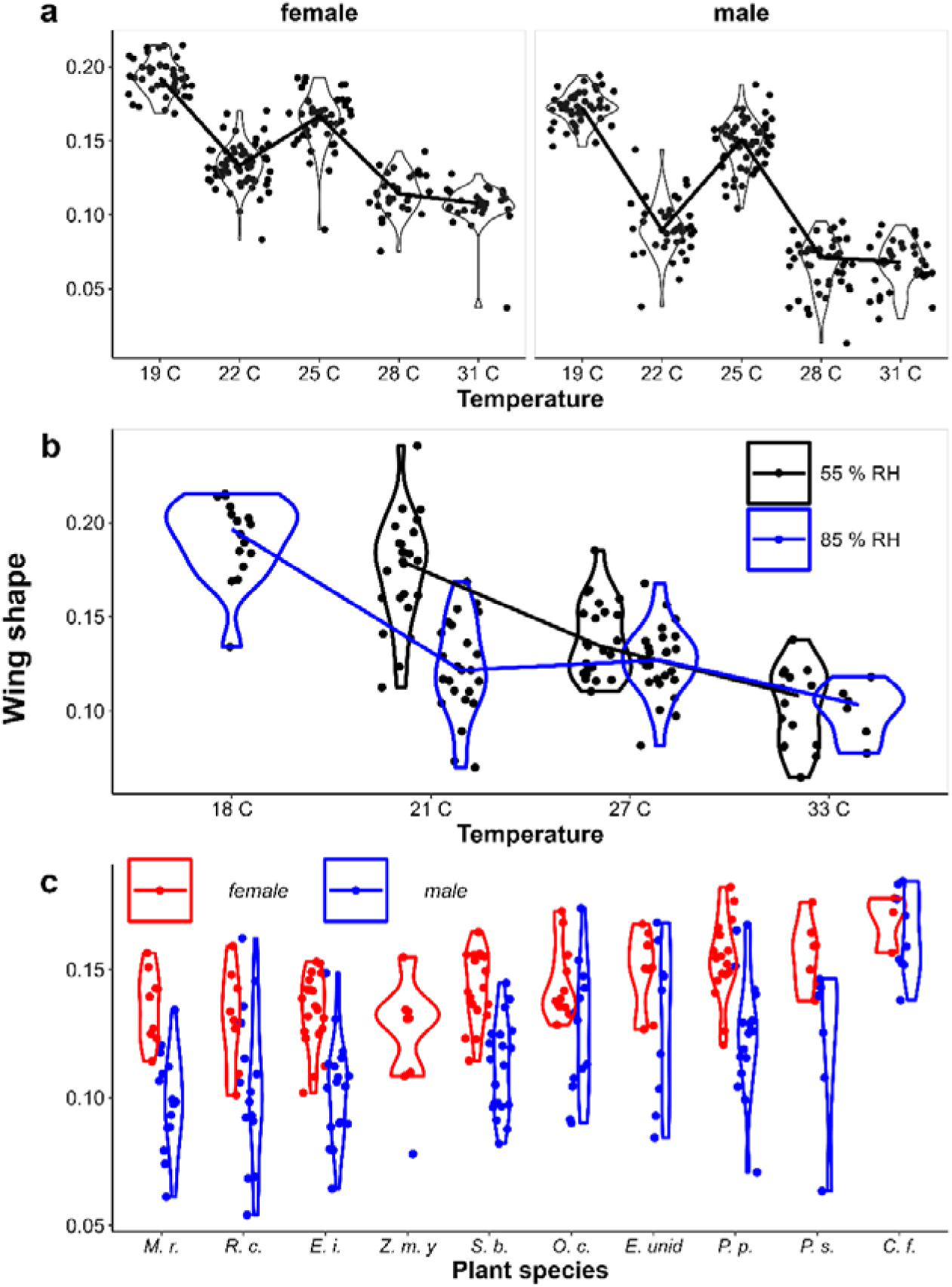
Provisional reaction norms of *M. leda* wing shape for a) the Temperature Experiment, b) the Temperature and Humidity Experiment, and c) the Host-Plant Experiment. Note that for 3b the data for high humidity for temperatures 21° and 27°C are from a pilot experiment starting from 2^nd^ instar caterpillars and sex was not recorded. For 3c only plants with N >7 were included and plants were sorted from small to large average eyespot size. Full plant species names are given in Table 1.

### Growth rate influencing eyespot size

While at higher temperatures larval growth was faster and adults tended to show more wet-season phenotypes (larger eyespots) in both the Ghanaian population (Figure 5a) and in the Indian Population (Figure 5b), there were only two cases of a significant relationship within temperature treatments in the Ghanaian population (Figure 5a, Table 1), and one in the Indian population (Figure 5b, Table 1). The adjusted R^2^ for these relationships tended to be very low (Table 1). Moreover, at intermediate temperatures, there were clear differences in adult phenotype between temperature treatments that produced a similar range of growth rates (22° and 25°C in Figure 3a, and 21° and 27°C in Figure 3b). Within host-plant species, growth rate was related to eyespot size in males (significant in one species of host plant, but the adjusted R^2^ was low), but not in females (Figure 5c, Table 1).

**Figure 5.**
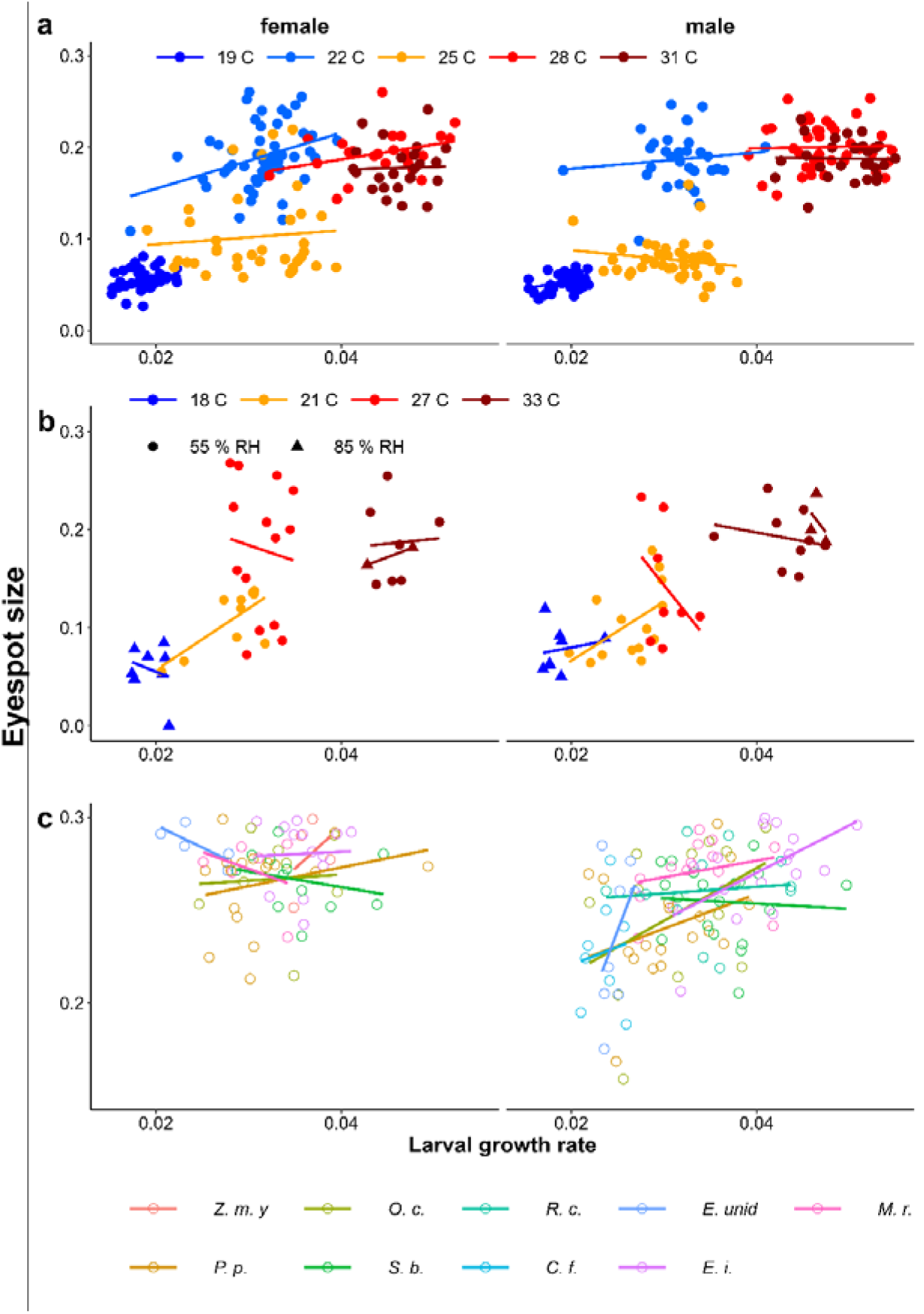
The effect of larval growth rate on eyespot size with a) the Temperature Experiment with the Ghanaian population, b) the Temperature and Humidity Experiment with the Indian population, and c) the Host-Plant Experiment with the Indian population. See Table 1 for sample sizes, statistical results, and full names of host plants.

**Figure 6.**
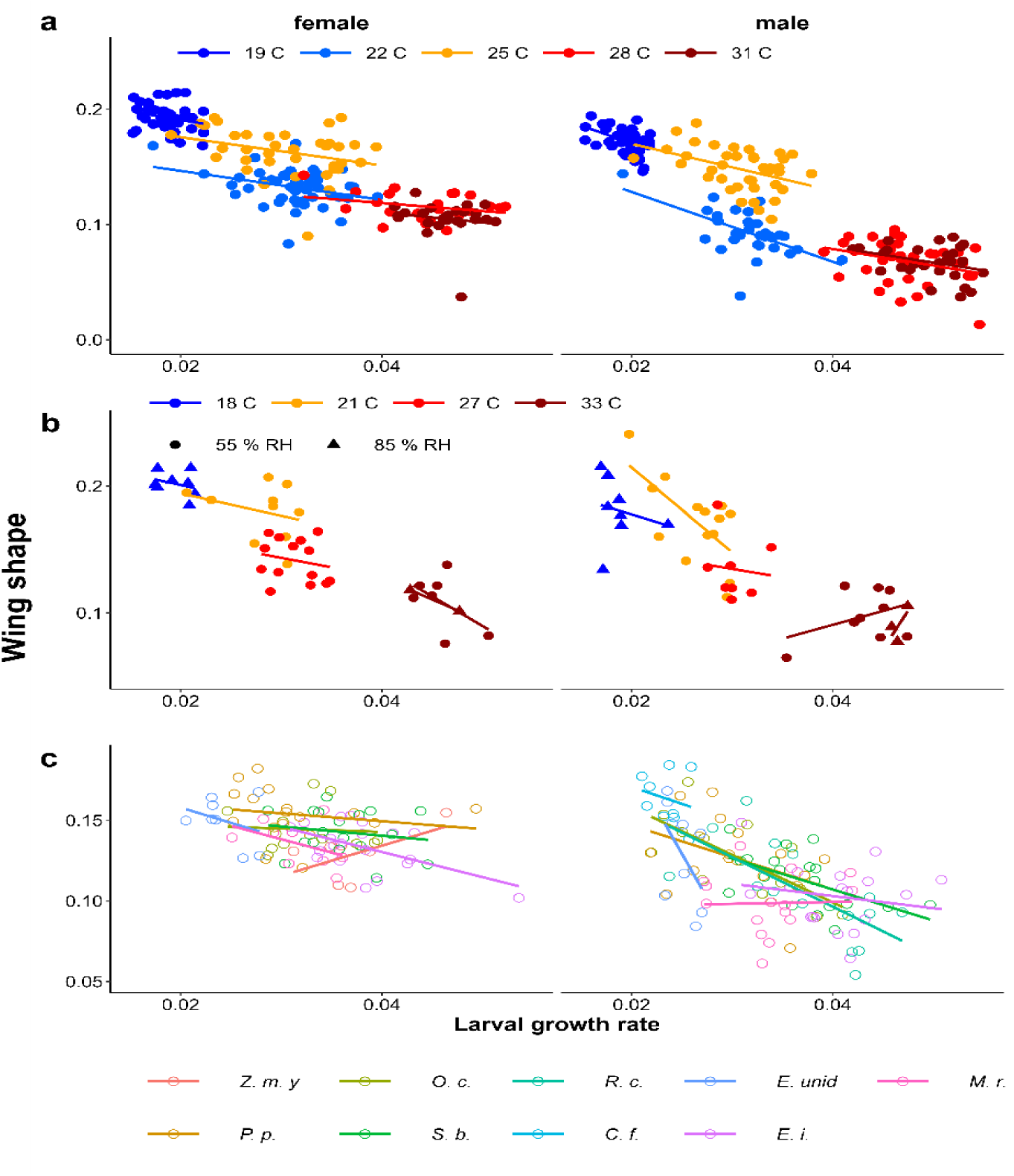
Scatter plots of larval growth rate and wing shape for a) the Temperature experiment (Ghana population), b) the Temperature and Humidity Experiment (Indian population), and c) the Host-Plant Experiment (Indian Population). See Table 2 for sample sizes and statistical results and Table 1 for full names of host plants.

### Growth rate influencing wing shape

In all experiments, lower larval growth rate was associated with more falcate wings in both sexes across and within treatments (Figure 4, Table 2). However, at intermediate temperatures, there were clear differences in adult phenotype between temperature treatments that produced a similar range of growth rates (22° and 25°C in Figure 4a, and 21° and 27°C in Figure 4b), and adjusted R^2^ tended to be quite low (Table 2). The relationships appeared to be stronger among males than among females, particularly for host plants (Figure 4c). Males tended to have less falcate wings when they were wet-season forms than females, so that the range of wing shapes was wider in males (most clearly seen in Figures 4a & c).

**Table 2.** Results of linear regression for wing shape. See Table 1 for abbreviations.

| Exp. | Temp | Humidity | Plant | sex | N | Estimate | S. E. | Statistic | p-value | R<sup>2</sup> | Adj. R<sup>2</sup> |
| --- | --- | --- | --- | --- | --- | --- | --- | --- | --- | --- | --- |
| Temperature Experiment, Ghanaian population |  |  |  |  |  |  |  |  |  |  |  |
|  | 31 C | 75% | Z. m. y | female | 25 | -0.94 | 0.96 | -0.98 | 0.34 | 0.04 | 0.00 |
|  | 28 C |  |  |  | 23 | -0.70 | 0.48 | -1.44 | 0.16 | 0.09 | 0.05 |
|  | 25 C |  |  |  | 37 | -1.22 | 0.62 | -1.99 | 0.05 | 0.10 | 0.08 |
|  | 22 C |  |  |  | 52 | -1.26 | 0.51 | -2.48 | **0.02** | 0.11 | 0.09 |
|  | 19 C |  |  |  | 39 | -1.55 | 0.93 | -1.67 | 0.10 | 0.07 | 0.05 |
|  | 31 C |  |  | male | 31 | -1.01 | 0.61 | -1.66 | 0.11 | 0.09 | 0.05 |
|  | 28 C |  |  |  | 41 | -1.15 | 0.62 | -1.84 | 0.07 | 0.08 | 0.06 |
|  | 25 C |  |  |  | 40 | -2.03 | 0.70 | -2.91 | **0.01** | 0.18 | 0.16 |
|  | 22 C |  |  |  | 32 | -3.03 | 0.78 | -3.87 | **0.00** | 0.33 | 0.31 |
|  | 19 C |  |  |  | 44 | -3.02 | 0.88 | -3.43 | **0.00** | 0.22 | 0.20 |
| Temperature and Humidity Experiment, Indian population |  |  |  |  |  |  |  |  |  |  |  |
|  | 18 C | 85% | Z. m. y | female | 8 | -1.73 | 2.25 | -0.77 | 0.47 | 0.09 | -0.06 |
|  | 21 C | 55% |  |  | 10 | -1.78 | 2.06 | -0.86 | 0.41 | 0.08 | -0.03 |
|  | 27 C | 55% |  |  | 14 | -1.48 | 2.01 | -0.74 | 0.47 | 0.04 | -0.04 |
|  | 33 C | 55% |  |  | 7 | -4.66 | 3.53 | -1.32 | 0.24 | 0.26 | 0.11 |
|  | 18 C | 85% |  | male | 8 | -2.45 | 4.73 | -0.52 | 0.62 | 0.04 | -0.12 |
|  | 21 C | 55% |  |  | 14 | -6.70 | 2.23 | -3.01 | **0.01** | 0.43 | 0.38 |
|  | 27 C | 55% |  |  | 8 | -1.31 | 5.07 | -0.26 | 0.80 | 0.01 | -0.15 |
|  | 33 C | 55% |  |  | 9 | 2.18 | 1.99 | 1.10 | 0.31 | 0.15 | 0.03 |
|  | 33 C | 85% |  |  | 3 | 12.21 | 12.25 | 1.00 | 0.50 | 0.50 | 0.00 |
| Host Plant Experiment, Indian population |  |  |  |  |  |  |  |  |  |  |  |
|  | 24 C | 69% | Z. m. y | female | 7 | 1.86 | 1.26 | 1.48 | 0.20 | 0.30 | 0.16 |
|  |  |  | *P. p.* |  | 17 | -0.49 | 0.67 | -0.73 | 0.48 | 0.03 | -0.03 |
|  |  |  | *O. c.* |  | 12 | -0.20 | 0.97 | -0.21 | 0.84 | 0.00 | -0.10 |
|  |  |  | *S. b.* |  | 18 | -0.57 | 0.81 | -0.70 | 0.49 | 0.03 | -0.03 |
|  |  |  | *E. unid* |  | 8 | -1.90 | 2.27 | -0.83 | 0.44 | 0.10 | -0.05 |
|  |  |  | *E. i.* |  | 20 | -1.56 | 0.57 | -2.73 | **0.01** | 0.29 | 0.25 |
|  |  |  | *M. r.* |  | 9 | -1.66 | 1.30 | -1.28 | 0.24 | 0.19 | 0.07 |
|  |  |  | *P. p.* | male | 18 | -1.98 | 1.06 | -1.87 | 0.08 | 0.18 | 0.13 |
|  |  |  | *O. c.* |  | 13 | -2.95 | 0.84 | -3.51 | **0.00** | 0.53 | 0.48 |
|  |  |  | *S. b.* |  | 19 | -1.93 | 0.68 | -2.85 | **0.01** | 0.32 | 0.28 |
|  |  |  | *R. c.* |  | 15 | -3.07 | 0.91 | -3.38 | **0.00** | 0.47 | 0.43 |
|  |  |  | *C. f.* |  | 9 | -2.10 | 3.39 | -0.62 | 0.56 | 0.05 | -0.08 |
|  |  |  | *E. unid* |  | 9 | -10.98 | 6.72 | -1.63 | 0.15 | 0.28 | 0.17 |
|  |  |  | *E. i.* |  | 18 | -0.76 | 1.00 | -0.76 | 0.46 | 0.03 | -0.03 |
|  |  |  | *M. r.* |  | 15 | 0.12 | 1.18 | 0.10 | 0.92 | 0.00 | -0.08 |

Overall, eyespot size was rarely related to growth rate within treatments (Table 1), while wing shape was consistently (but still weakly) related to growth rate across all three experiments and in both sexes (Table 2). These effects were generally more pronounced in males than in females (Tables 1 and 2).

## DISCUSSION

We reared more than 800 larvae of *M. leda* under various temperature, humidity, and host-plant conditions. Overall, larvae reared at higher temperatures tended to produce more wet-season-form phenotypes (larger eyespots and less falcate wing tips), and host-plant species also affects adult phenotype, similar to what has been found in some *Bicyclus* butterflies. We found little evidence that larval growth rate mediates between environmental cues and eyespots (only some significant relationships for males in the Host-Plant Experiment and females in the Temperature Experiment), while growth rate was consistently related to wing shape within treatments. Notably, even when growth rate clearly affected adult phenotypes, within treatments, there were large differences in adult phenotype between treatments that produced a similar range of growth rates. These results suggest that there is no direct link between growth rate and eyespot size, while wing shape is in part, but not solely, determined by larval growth rate. Moreover, there are differences between the sexes.

*M. leda* larvae are more likely to develop into wet-season-form adults when reared at higher temperatures, and on host-plant species on which they grow faster. That higher temperatures induced wet-season phenotypes similar to distantly related satyrines (van Bergen et al. 2017) may be explained by the increases in temperature as the wet season is approaching and decreases in temperature at the end of the wet season (Halali et al. 2021; van Bergen & Oostra 2023). Temperature during larval development is thus likely to be predictive of the season the adult will experience. Using host-plant quality as a cue may be adaptive because towards the end of the wet season, plants will often be of lower quality, and plant quality can thus be a reliable predictive cue for larvae: low plant quality indicating that the dry season is approaching. Conversely, once the rains have started, caterpillars will on average experience high food quality on which they can achieve a faster growth rate. Since butterflies will probably start laying eggs at the beginning of the wet season, plant quality is probably a reliable cue during this period. Further studies may test whether the developmental plasticity for these two cues and two traits was inherited from a common ancestor or case of parallel evolution. (Bhardwaj et al. 2020).

This climate pattern is common in the tropics, but does not apply everywhere. Therefore, the response to temperature might vary between populations (Roskam & Brakefield 1996), similar to findings in *B. anynana* and *B. safitza* (de Jong et al. 2010; Nokelainen et al. 2018). While the Indian and Ghanaian populations responded qualitatively similarly to temperature in our experiments, a quantitative comparison among the populations would require rearing them simultaneously in a common garden experiment. Furthermore, the reaction norms should be considered as provisional. For example, possible effects of growth chamber or outdoor weather conditions cannot be excluded in our results. In particular, the strange shape of the temperature reaction norm for the Ghanaian population, and the effects of humidity need to be investigated further.

Our results suggest that different cues might in part be integrated through growth rate for wing shape, but not for eyespot size. Even for wing shape, larval growth does not appear to be solely responsible for determining adult phenotype: adjusted R^2^ was typically low, and at intermediate temperatures, growth rates were similar while adult phenotypes differed greatly among temperature treatments. Perhaps growth rate during a small section of the development time is the actual cue (a critical period, Kooi & Brakefield 1999). Since larval development may be slow during the first instars on certain plants but then accelerate in later instars, overall growth rate might be a poor predictor of adult phenotype if there is such a critical period. However, this would not explain the large differences in adult phenotype between treatments that produced nearly identical growth rates such as at 22°C and 25°C in the Temperature Experiment. Therefore, growth rate is probably not the determining factor affecting adult phenotype, even for wing shape.

It may be surprising that eyespot size and wing shape are regulated differently. This can lead to individuals with a mix of dry and wet-season phenotypic traits such as large eyespots combined with falcate wing tips. In other satyrines, multiple plastic traits tend to be linked, but there are also exceptions (Mateus et al. 2014; van Bergen et al. 2017). While such disintegration of traits may be expected to be disadvantageous, it could also contribute to the extensive variation among individuals (Ruiter & Brakefield 1994) that may hamper search-image formation in predators (Cook 2017; Forsman 2015; Forsman et al. 2015).

We found notable sex differences in the relationship between larval growth rate and adult phenotype. Perhaps females often have a maximum eyespot size when they are wet-season form, truncating their size distribution and thus limiting their potential response to growth rate, while in males eyespots tend to be smaller and to cover a wider range of sizes (see Figure 3c). For wing shape, the effects of growth rate also tended to be more pronounced in males than in females. Specifically, males tended to have even less falcate wings when they were wet-season forms than females. This indicates some subtle sexual dimorphism in wing shape in wet-season forms that may be connected to sex-biased movement patterns (Berwaerts et al. 2002) or sexual selection (Kemp 2002; Molleman et al. 2020a).

In conclusion, we found that cue use of *M. leda* is similar to that of distantly related satyrines. The mechanism of seasonal dimorphism in *M. leda* can be understood as developmental plasticity using temperature and certain aspects of host-plant quality as cues, where larvae reared at higher temperatures and on host plants on which they can grow faster tend to show more wet-season phenotypes (larger eyespots and less falcate wing tips). This cue use appears mediated in part by larval growth rate for wing shape, but not for eyespot size.

## Acknowledgments

We thank Kwaku Aduse-Poku for help in obtaining the population from Ghana. Funding for the work was provided by the University of Cambridge, the Indian Institute of Science Education and Research Thiruvananthapuram, and the Naradowe Centrum Nauki (National Science Centre, Poland) grant 2021/43/B/NZ8/00966.

## Author Statement

EMM performed the Temperature Experiment, united the data sets and performed preliminary data analysis, SH performed the Temperature and Humidity Experiment and performed preliminary data analysis, FM performed the Host-Plant Experiment supervised the work of SH, performed the final data visualization and analysis, and wrote the first draft of the manuscript. VO supervised the work of EMM and advised on data analysis. EvB wrote the ImageJ code for measuring wing traits and co-supervised EMM. DH analyzed the images from the Host-Plant Experiment. PMB co-supervised EMM. UK supervised SH. All co-authors contributed to the writing of the manuscript.

