## Appendix 1 for "Larval growth rate affects wing shape more than eyespot size in the seasonally polyphenic butterfly *Melanitis leda*"

Results of follow-up experiment to verify unexpected non-linear reaction norm in the Temperature Experiment.

Methods similar to Temperature Experiment (see main text).

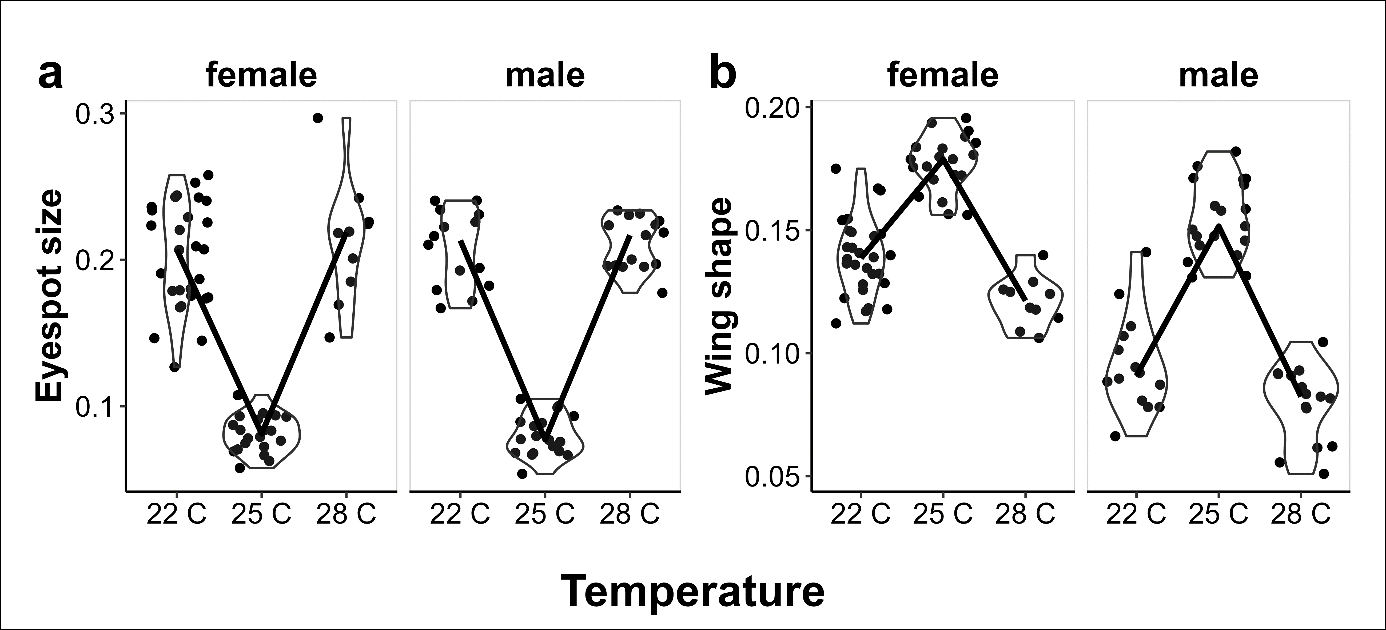

Figure A1.1. Provisional reaction norms of *M. leda* for a) Eyespot size, and b) wing shape.

The results are similar to those of the main experiment (see Figure 3 in main text): a non-linear effect of temperature on adult phenotype.
